# Karyotype Evolution in Response to Chemoradiotherapy and Upon Recurrence of Esophageal Adenocarcinomas

**DOI:** 10.1101/2024.02.28.582275

**Authors:** K. van der Sluis, J. W. van Sandick, W. J. Koemans, T. van den Bosch, A. Broeks, D. Peters, I. M. Seignette, C. R. Rausch, E. van Dijk, P. Snaebjornsson, J. G. van den Berg, N. C. T. van Grieken, B. Ylstra, B. Carvalho, D. M. Miedema, L. L. Kodach

**Affiliations:** Department of Surgical Oncology, Netherlands Cancer Institute, Amsterdam, The Netherlands; Amsterdam UMC location University of Amsterdam, Cancer Center Amsterdam & Amsterdam Gastroenterology Endocrinology Metabolism, Center for Experimental and Molecular Medicine, Amsterdam, The Netherlands; Oncode Institute, Amsterdam, The Netherlands; Core Facility Molecular Pathology and Biobanking, Netherlands Cancer Institute, Amsterdam, The Netherlands; Department of Pathology, Netherlands Cancer Institute, Amsterdam, The Netherlands; Faculty of Medicine, University of Iceland, Reykjavik, Iceland; Department of Pathology, Cancer Center Amsterdam, Amsterdam UMC, Vrije Universiteit Amsterdam, Amsterdam, The Netherlands

## Abstract

The genome of esophageal adenocarcinoma (EAC) is highly unstable and might evolve over time. Here, we track karyotype evolution in EACs in response to treatment and upon recurrence through multi-region and longitudinal analysis. To this end, we introduce L-PAC, a bio-informatics technique that allows inference of absolute copy number aberrations (CNA) of low-purity samples by leveraging information of high-purity samples from the same cancer. Quantitative analysis of matched absolute CNAs reveals that the amount of karyotype evolution induced by chemoradiotherapy (CRT) is predictive for early recurrence and depends on the initial level of karyotype intra-tumor heterogeneity. We observe that CNAs acquired in response to CRT are partially reversed back to the initial state upon recurrence. CRT hence alters the fitness landscape to which tumors can adjust by adapting their karyotype. Together, our results indicate that karyotype plasticity contributes to therapy resistance of EACs.

## Introduction

Aneuploidy, an abnormal number of chromosomes in a cell, occurs in around 90% of cancers (1, 2). Although aneuploidy is an almost universal feature of cancer, cancer (sub-)types have a characteristic karyotype with unique copy number alterations (CNAs). For example, squamous cell cancers of the lung frequently have a loss of chromosome 13, while most colorectal cancers have more than two copies of chromosome 13 (1). The context dictates which specific karyotype is selected and becomes clonal (3, 4). Indeed, the rationale to treat cancers with systemic- and/or radiotherapy is to alter the selective pressure acting on cancer cells. Some cancer cells, however, adapt to the selective pressure imposed by therapy and cause therapy resistance (5). This adaptation of the cancer cell population to therapy might be achieved through selection of specific karyotypes (6, 7). Recent work by Lukow *et al*. (8) and Ipolito *et al*. (9) demonstrated that such adaptation of the cancer cell karyotype upon treatment occurs *in vitro*. Whether a similar karyotype evolution occurs in patients during treatment has not yet been studied in detail.

Patients with locally advanced esophageal adenocarcinoma (EAC) are treated with neo-adjuvant chemoradiotherapy (nCRT) or perioperative chemotherapy. However, 82% of EACs have an incomplete pathological response to nCRT (10). The prognosis of patients with locally advanced EAC is indeed dismal, with a 5-year survival rate of 24%. A pressing clinical concern is hence understanding why EACs are refractory to therapy.

Previous work has shown that the genome of EACs evolves in response to treatment and upon disease progression. A longitudinal study of EAC including samples from autopsies suggested ongoing selection during the progression of disease (12). This is further supported by reports of “bottlenecking”, the selection of one or a few clones, in response to treatment of EACs with chemotherapy, in particular in responders (13). Moreover, a recent study suggested more sub-clonal CNAs in non-responding EAC upon treatment (14). Hence, previous work has demonstrated that EACs evolve in response to treatment (15) and have a high level of chromosomal instability (CIN) (16). Yet, if and how CIN contributes to the evolution of EAC remains unclear.

The evolution of malignant cells occurs in interaction with a complex ecologic system of stroma, vasculature and immune infiltrate (17). In particular the immune tumor microenvironment (iTME) has been shown to impose a strong selective pressure during tumor evolution (18). This study showed that immune evasion occurred via neo-antigen depletion through loss of heterozygosity. The level of CNA mediated immune-editing was associated with the density of immune infiltration. Importantly, adaptive karyotype changes in response to treatment with immune checkpoint inhibitors (ICI), such as neo-antigen elimination via chromosome losses, might contribute to acquired resistance for ICI treatment in lung cancer (19). As ICI have recently been shown to be effective in EAC treatment for a subset of patients (20), analysis of how the immune infiltrate shapes tumor evolution may allow identification of those patients more likely to benefit from adjuvant ICI therapy.

Here, we investigated the evolution of karyotypes in EACs in response to treatment with nCRT and upon tumor recurrence by longitudinal and multi-region measurements of CNAs. We first introduce **L**ow **P**urity inference of **A**bsolute **C**NAs (L-PAC), a bio-informatics technique that leverages the information of high-purity samples to improve the inference of absolute CNAs for low-purity samples from the same patient. The multi-region and longitudinal absolute CNA measurements are next quantitatively compared to track karyotype evolution in EAC in response to nCRT, upon recurrence and in the context of the iTME.

## Results

The evolution of EAC karyotypes in response to nCRT and upon recurrence was analyzed by performing shallow whole genome sequencing (sWGS) on 170 matched samples acquired before nCRT (*preCRT)*, from the surgical resection after nCRT (*postCRT)* and upon *recurrence* from 24 patients with EAC (Figure 1A). All 24 patients were treated with nCRT and had a recurring disease, resulting in a median overall survival of 20 months (Interquartile Range (IQR) 7-29) (Table S1). A median of 7 (IQR 6-9) tumor regions were analyzed per patient, with at least one sample analyzed per time point for each patient.

**Figure 1.**
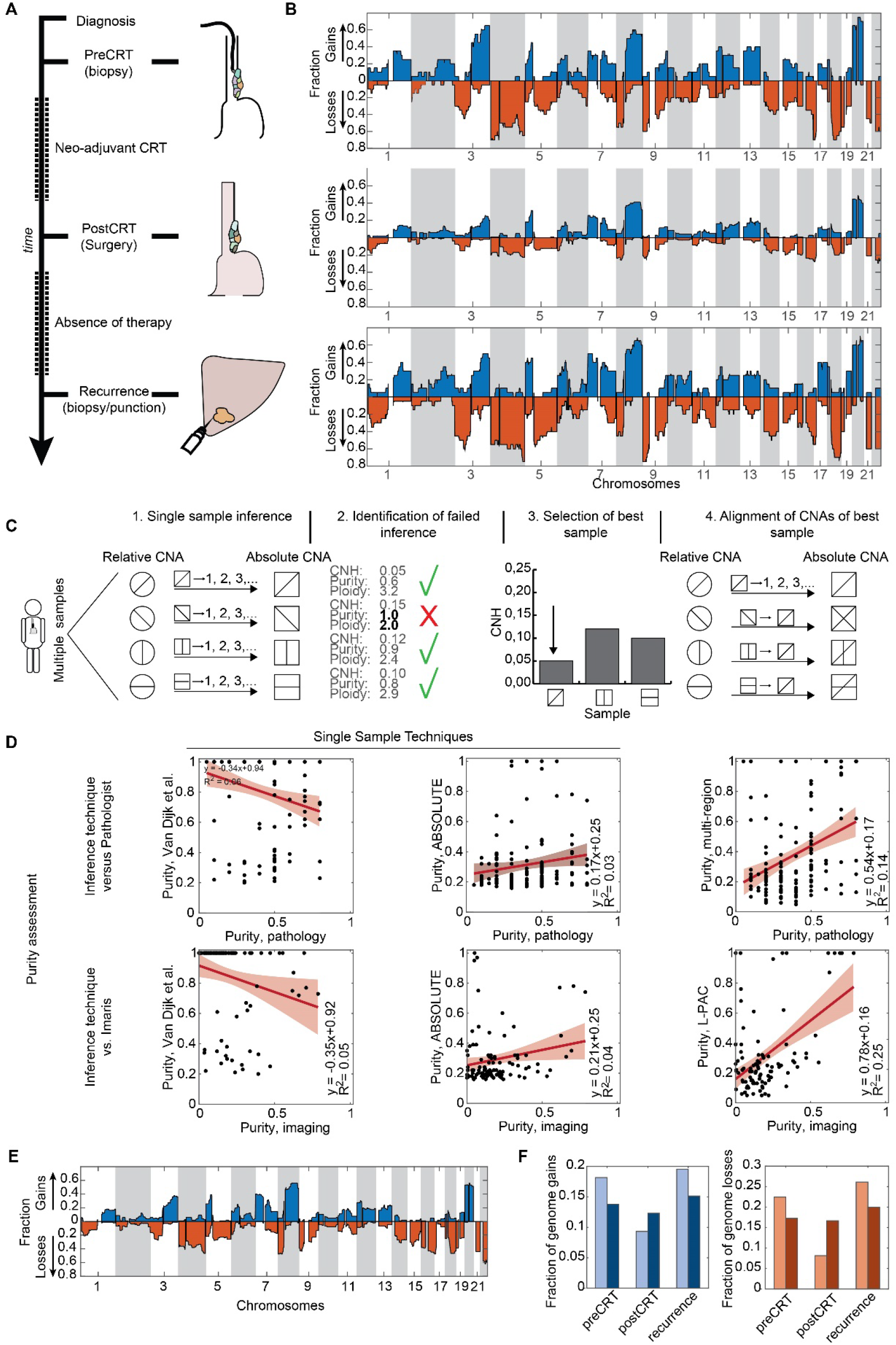
Longitudinal measurement and inference of absolute CNAs. **A**, Study design. Longitudinal sampling of EACs: biopsy taken before nCRT (*preCRT*), multi-region sampling from resected tumor after nCRT (*postCRT*) and biopsy taken of tumor after recurrence (*recurrence*). **B**, Fraction of samples with gains and losses at each position in the genome based on relative CNAs for samples taken *preCRT* (upper panel), *postCRT* (middle panel) and *recurrence* (lower panel). **C**, L-PAC workflow for inference of absolute CNAs from multiple samples from a tumor. **D**, Comparison of inferred purity from CNAs to purity assessed by pathologist (upper panels) and purity reported by HALO imaging software (lower panels). Linear regression with corresponding p values are reported. **E**, Fraction of gains and losses in *postCRT* samples from absolute CNAs derived by L-PAC. **F**, histogram showings fraction of genome gained (left panel) and lost (right panel) per time point for relative CNAs (light) and absolute CNAs inferred by L-PAC (dark).

GISTIC analysis of focal CNAs indicated gains on chromosome 1p36.2, 2p16.1, 2q32.1, 3q26.2, 5p13.2, 6p21.1, 7q21.2, 11q13.3, 12p12.1 and 13q22.3 and losses on chromosomes 1p35.3, 2p22.3, 2q37.2, 3p21.1, 4p16.3, 4q35.1, 6p25.1, 6q15, 8p21.1 and 9q21.3 (Figure S1), in line with previous reports of common focal CNAs in EACs (7, 15, 21, 22). Furthermore, analysis of the genome wide relative copy numbers reaffirmed previous findings of large CNAs within the tumor genome, e.g., gains of chromosome arms 3q and 8q (Figure 1B).

Analysis of each time point separately revealed a reduced frequency of CNAs in *postCRT* samples (Figure 1B). The apparent decrease in CNAs could be an artefact arising from the diminished tumor cell percentage after nCRT. Indeed, our pathologist reported a lower median purity of *postCRT* samples (30% (IQR 19-45) compared to both *preCRT* (50% (IQR 30-60)) and *recurrence* samples (50% (IQR 49-53)). We employed ABSOLUTE to infer the tumor-cell only, or absolute, CNAs from relative CNAs (23). However, ABSOLUTE yielded a, for EACs, unrealistic low percentage of CNAs. Indeed, methods such as ABSOLUTE that infer absolute CNAs require a sufficiently high tumor cell purity and are therefore not well suited for the analysis of the current dataset (24).

### Multi-region inference of absolute CNAs

To facilitate the accurate assessment of CNAs in low purity samples, which is crucial for tracking karyotype evolution in EAC, we introduced a multi-region CNA inference method (L-PAC) that leverages the combined information from multiple available samples from the same tumor. In brief, L-PAC works as follows (see STAR methods for details): first, single-sample inference of absolute CNAs for each sample was performed through a grid-search of candidate ploidies and purities as described in Van Dijk *et al*. (25). In this first grid search, the average distance of candidate absolute CNAs to the closest integer value is minimized. Second, samples for which this single-sample inference failed were identified by an inferred tumor cell purity of 1 and a ploidy below 2.5 (see STAR methods for the mathematical argumentation). Third, intra-tumor heterogeneity (ITH) could hamper an accurate identification of absolute CNAs. Hence, we identified for each tumor the best sample β for which single-sample inference did not fail, as the sample with the lowest CNH (25). Fourth, we leveraged the inferred absolute CNAs of the best sample β to infer the absolute CNAs of the other samples of a tumor through a second grid search over purities and ploidies (Figure 1C). In this second grid search, the distance of candidate absolute CNAs to the absolute CNAs of sample β was minimized.

The performance of CNA inference by L-PAC was assessed by comparing the inferred purities to independent purity observations by a pathologist and HALO imaging software. Purity inferred by L-PAC was in reasonable concordance with the independent purity assessments (Figure 1D). Importantly, L-PAC performed better compared to the single-sample inference by ABSOLUTE (23) or the method used in Van Dijk et al. (25) (Figure 1D). For three patients no best sample β could be determined with L-PAC. In addition, one patient had a microsatellite instable tumor without any detectable CNAs. Therefore, these four patients were excluded from further analysis. Tracking of karyotype evolution was hence performed on a total of 149 samples from 20 patients in this study.

### Karyotypes evolve over time

To study karyotype evolution in EACs treated with nCRT, the difference in genome wide absolute CNAs between samples was quantified. Quantification was performed by measuring the genome average of the Manhattan distance between CNAs measured in two samples (dCNAs) (Figure 2A). We observed a significantly larger dCNA upon treatment (measured between *preCRT* and *postCRT* samples) and recurrence (measured between *postCRT* and *recurrence* samples) compared to the spatial ITH in *postCRT* samples (Figure 2B and 2C). Hence, although copy number ITH was observed between different regions evaluated at the same point in time, the largest variations in CNAs occurred over time, in particular upon recurrence (Figure 2D). Furthermore, hierarchical clustering of samples per patient based on pair-wise dCNAs also indicates that *postCRT* samples cluster together for most tumors (Figure 2E). Note that for several patients (patient 3, 8, 16 and 20) the *preCRT* and *recurrence* samples are more similar to each other than to any of the *postCRT* samples. We next addressed the biology driving karyotype evolution in response to CRT and upon recurrence.

**Figure 2.**
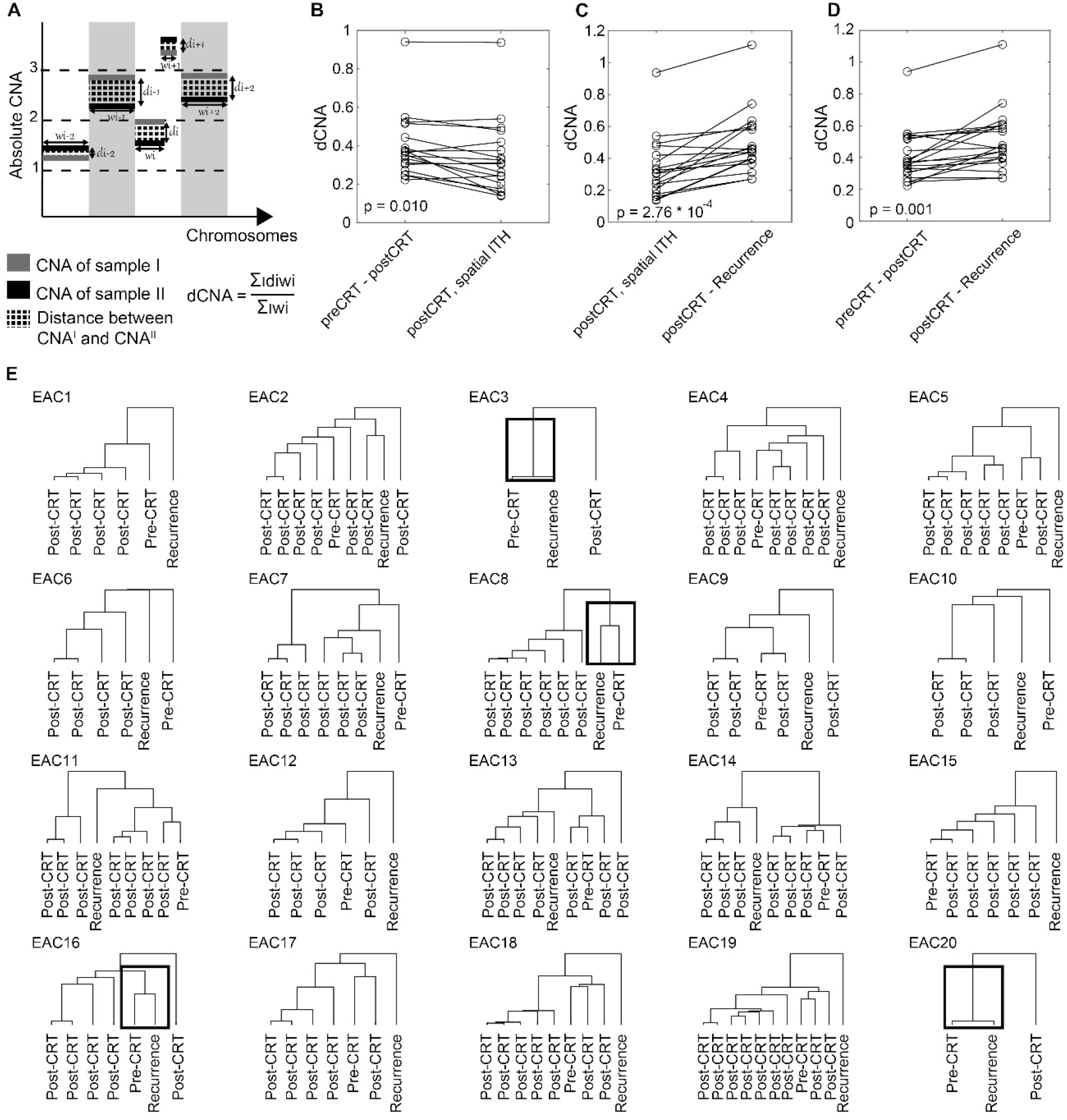
Quantitative analysis of karyotype evolution. **A**, Schematic representation of the average distance in copy number alterations (dCNA) between absolute CNAs from two samples. **B**, Paired analysis of dCNA measured between *preCRT* and *postCRT* compared to the average dCNA between multi-region samples *postCRT*. The p value is derived from the Wilcoxon signed rank test. **C**, Paired analysis of dCNA measured between *postCRT* and *recurrence* compared to the average dCNA between multi-region samples *postCRT*. The p value is derived from the Wilcoxon signed rank test. **D**, Paired analysis of dCNA measured between *preCRT* and *postCRT* compared to dCNA measured between *postCRT* and *recurrence*. The p value is derived from the Wilcoxon signed rank test. **E**, Clonal trees per patient obtained from hierarchical clustering of dCNA between samples. Tumors EAC3, EAC8, EAC16 and EAC20 are marked based on clustering of the *preCRT* sample with the *recurrence* sample.

### Karyotype evolution in response to CRT

First, we focused on the karyotype evolution induced by the pressure of nCRT. We observed that the amount of karyotype evolution in response to nCRT associated with the level of karyotype ITH (CNH) present before treatment (Figure S3A). ITH is a predictor of tumor progression, therapy resistance and a poor prognosis precisely because it fuels tumor evolution (17, 26). We thus reasoned that dCNA between the *pre-* and *postCRT* might be related to the future course of disease. As all 20 patients in the present study had a disease recurrence, we focused on the relation between dCNA and timing of recurrence. We observed that tumors with a relatively large value of dCNA in response to nCRT had a recurrence within 12 months, whilst tumors with a relatively low dCNA in response to nCRT recurred later (Figure 3A). Hence, our data indicates that a tumor with a karyotype that evolves in response to nCRT recurs fast.

**Figure 3.**
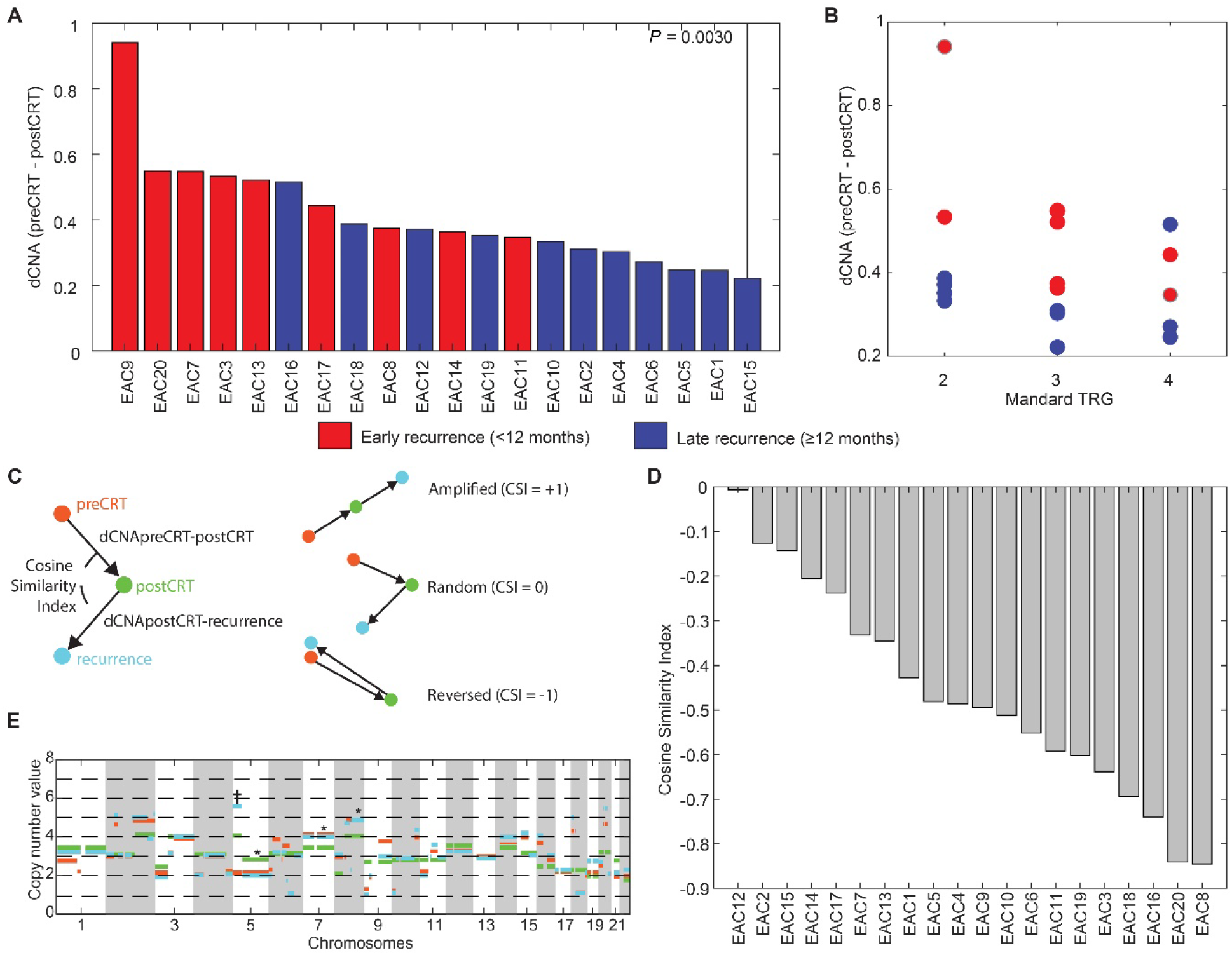
Karyotype evolution in response to chemoradiotherapy is associated with early recurrence and is partially reversed upon recurrence. **A**, Waterfall plot of copy number alteration distances (dCNA) between *preCRT* and *postCRT* per tumor. The color indicates early (<12 months, red) or late recurrences (>12 months, blue). dCNA between early and late recurrences are compared with the Wilcoxon rank sum test. **B**, dCNA between *preCRT* and *postCRT* versus the Mandard tumor regression grade (TRG) (Spearman’s rank correlation= -0.35; p = 0.13). Mandard TRG is not associated with early recurrence (p = 0.33, one-way ANOVA). **C**, Schematic representation for the definition of the cosine similarity index (CSI). **D**, CSI per tumor. **E**, Example of karyotype evolution in EAC3. Markers highlight reversed (*) and amplified (†) alterations.

The efficacy of nCRT is clinically assessed by the Mandard Tumor Response Grade (TRG), which categorizes the amount of tumor regression (27). Interestingly, no relation between the Mandard TRG and time of recurrence is observed (one-way ANOVA p = 0.33; Figure 3B). However, the Mandard TRG is inversely related to dCNA, with regressing tumors due to nCRT (i.e., low Mandard TRG) associated with more changes in the tumor genome (i.e., high dCNA) (Spearman’s rank correlation = -0.35; p = 0.13). Importantly, we observed that independent of the Mandard TRG, high dCNA is associated with swift tumor recurrence (Figure 3B). Thus, for patients with incomplete response and recurrent disease, the amount of karyotype evolution induced by nCRT is more predictive for the progression free interval than the Mandard TRG.

### Karyotype evolution upon recurrence

The tumor karyotype continues to evolve upon recurrence (Figure 2D). The level of karyotype ITH (CNH) *postCRT* was associated with the amount of karyotype evolution during *recurrence*, analogous to our observation for karyotype evolution in response to nCRT (Figure S3B). Hence, CNH is a predictor of karyotype evolution in EAC.

As noted above, we observed that *recurrence* samples of patients 3, 8, 16 and 20 clustered together with the *preCRT* samples (Figure 2E). This suggests that, at least for a subset of patients, CNAs accumulated in the genome in response to nCRT and recurrence are not incremental or random, but rather partially reversible. To quantify the relative direction of the subsequent CNAs we determined the cosine similarity between the CNA changes in response to nCRT and CNA changes during recurrence (Figure 3C). Strikingly, we found that for all patients the cosine similarity of CNAs is negative, implying that part of the CNAs acquired in response to nCRT are reversed during recurrence (Figure 3D and 3E). In the period prior to recurrence, tumors evolved without pressure from nCRT. As such, the selective pressure during recurrence is likely more akin to the selective pressure prior to nCRT *(preCRT)*, than resulting from nCRT. We conjecture that the relative similarity of the karyotypes before nCRT and at recurrence, compared to immediately after nCRT, is explained by selective pressure imposed by nCRT on karyotypes and the ability of the tumor karyotype to adapt to changing selective pressure.

### Impact of the immune tumor microenvironment on recurrence

Finally, we explored the role of the iTME on tumor evolution during recurrence through multi-region analysis of *postCRT* samples with multiplex immunofluorescence. A wide range of immune infiltration was found between patient (Figure 4A) and within patients (Figure 4B). We analyzed if the local level of immune infiltrate of a sample correlates to the probability of the recurrence originated from the sample. The probability of seeding the recurrence for each *postCRT* sample was determined based on the similarity of the *postCRT* sample karyotype to the karyotype of the *recurrence* (see STAR Methods). Analyzing all patients combined, we did not find a significant relation between similarity to recurrence of a *postCRT* region and the local density of the immune infiltrate in the *postCRT* region (Figure S4). However, in patients with early recurrence (within 12 months), we observe a significant correlation for the CD8/CD3 and the FoxP3/CD8 ratio (Figure 4C and 4D).

**Figure 4.**
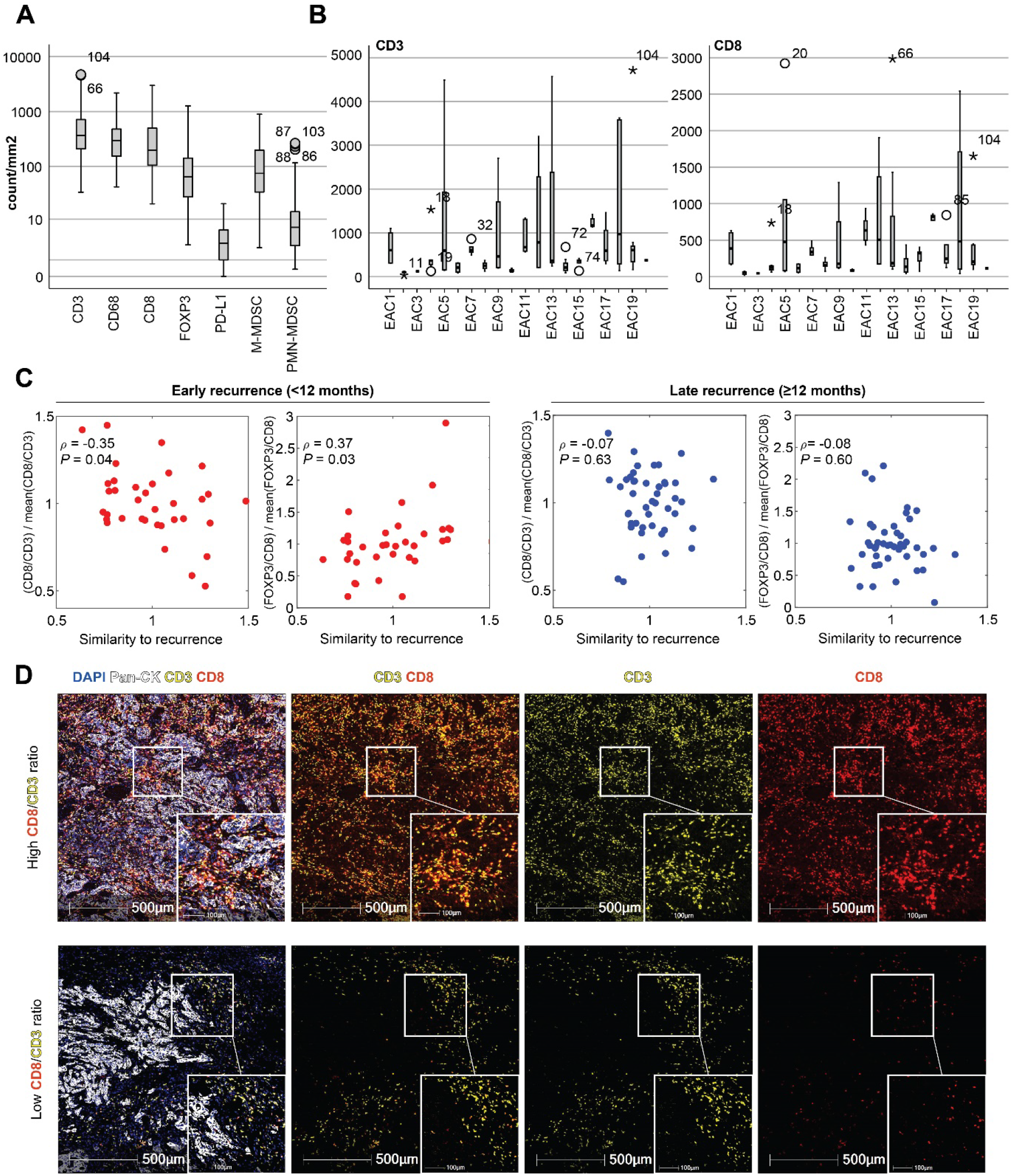
Impact of the iTME on tumor evolution. **A**, Density of cells with positive staining within all *postCRT* samples. **B**, Density of CD3 (left panel) and CD8 (right panel) positive cells within *postCRT* samples per tumor of all patients. **C**, Scatterplots of CD8/CD3 ratio and FOXP3/CD8 ratio in *postCRT* samples versus the similarity to recurrence of the samples, for early (<12 months) and late (>12 months) recurrences. The Spearman’s rank correlation coefficients with corresponding p values are reported. **D**, Multiplex immunofluorescence imaging of a sample from patient 16 (upper row) and a sample from patient 4 (lower row) illustrates the observed variability in CD8/CD3 density.

## Discussion

In this work we tracked karyotype evolution in response to nCRT and upon recurrence in EAC. While previous works tracked the clonal evolution of EACs in detail (12, 13, 15), relatively little is known of the karyotype evolution of EAC, in particular in response to nCRT. Our results demonstrate that the tumor karyotype of EACs shows plasticity in response to nCRT and upon recurrence. We found that the amount of change in the tumor karyotype can be predictive for early and late recurrences and was depended on the initial level of karyotype heterogeneity as well as the iTME. Moreover, our results suggest that the karyotype appears to adapt to the selective pressure imposed by nCRT, and that this adaptation is reversible.

To longitudinally track karyotype evolution, we developed the L-PAC for the inference of absolute CNAs. Biopsies and post-treatment samples typically have a low percentage of tumor cells, hampering the inference of absolute CNAs from relative CNAs. By leveraging information from high-purity samples from the same tumor we could improve the inference accuracy of low-purity samples compared to single-sample inference techniques. Inference of absolute CNA is notoriously difficult (28), and previous methods have been developed that leverage the full information from multiple samples of a tumor to determine absolute CNAs (29, 30). We designed L-PAC specifically to improve the inference of absolute CNAs for low-purity samples. A limitation of L-PAC is that it ignores sub-clonal whole-genome doublings. The applicability of L-PAC could therefore be limited for tumor types where sub-clonal whole-genome doubling is a prevalent event (31).

ITH is a predictor of poor prognosis in cancer (32-34). The biological interpretation of this clinical observation is that diverse tumor cell populations are well able to adapt and evolve, e.g., under pressure from therapy. We observed that indeed the initial level of karyotype ITH correlated to the amount of change in CNAs when longitudinally tracking karyotypes. Moreover, an evolving tumor karyotype in response to nCRT was associated with early disease recurrence, independent of the amount of tumor regression as quantified by the Mandard TRG. These results suggest that a large change in the tumor karyotype after treatment reflects the tumor successfully passing through a selection bottleneck imposed by nCRT, in line with previous reports of bottlenecking of EACs upon treatment (13). All patients analyzed in the current work had recurrence of disease. Future work is needed to explore how karyotypes change upon treatment in patients with incomplete response to nCRT, but without recurrent disease.

The selection imposed by nCRT is only temporal and fades after withdrawal of therapy. Therefore, the selective pressure acting on a tumor before and after treatment are likely more similar than the selective pressure in response to treatment. The longitudinal karyotype measurements before nCRT, (immediately) after nCRT and months later at recurrence allowed us to address if tumor karyotypes of human *in situ* cancers adapt to selective pressure from therapy, as has been reported to occur *in vitro* (8, 9). However, the selective pressure from therapy works in concert with selective pressures arising from the unique tumor biology of each tumor. Moreover, in addition to specifically selected CNAs the karyotype can also consist of passenger CNAs. We reasoned that, nevertheless, if tumor unique changes in karyotype that emerged in response to treatment were (partially) reversed over time when treatment was withdrawn, this would be an indication of nCRT imposing selection on the karyotype, and the adaptability of the tumor karyotype to changing selective pressures. Strikingly, we found that for all 20 patients analyzed karyotype changes acquired in response to treatment are partially reversed after treatment.

The iTME is a second important factor that, in addition to nCRT, shapes tumor evolution (32, 36). For example, spatial variations of immune infiltrate within a tumor were observed to contribute to the clonal evolution in lung adenocarcinoma (37). Our results of multi-region spatial iTME in *postCRT* samples indicated that enrichment of the cytotoxic CD8-positive compartment and decrease in regulatory FOXP3-positive compartment appears to suppress EAC recurrence in patients with early recurrence. Our results are in line with the anti-tumor function of CD8-positive cells (38, 39).

To conclude, we observed that karyotypes of EACs evolve in response to nCRT and upon recurrence. The observed plasticity of CNAs over time suggests that karyotype evolution contributes to adaptation of cancer cells to the selective pressure imposed by nCRT in EACs and thus plays a role in therapy resistance and disease recurrence. Moreover, the amount of karyotype change in response to nCRT could be a potential biomarkers to inform on the timing of future recurrence. Together, our results highlight the fundamental biology and potential clinical relevance of karyotype evolution in EACs.

## Methods

### Key Resources Table

| REAGENT or RESOURCE | SOURCE | IDENTIFIER |
| --- | --- | --- |
| Antibodies |  |  |
| Anti-CD68 | DAKO | Clone KP1, Cat M0814 |
| Anti-CD8 | DAKO | Clone C8/144B, Cat M7103 |
| Anti-FoxP3 | Abcam | clone 236A/E7, Cat ab20034 |
| Anti-CD3 | Thermo-Scientific | Clone SP7, Cat RM-9107-S |
| Anti-PanCK | Thermo-Scientific | Clone AE1/AE3, cat MS343P |
| Anti-CD20 | DAKO | Clone L26, Cat M0755 |
| Anti-CD11b | Cell Signaling | Clone D6X1N, Cat CST49420s |
| Anti-CD15 | Becton Dickinson | Clone MMA, Cat 559045 |
| Anti-CD14 | Abcam | Clone EPR3653, Cat ab133335 |
| Anti-HLA-DR | Abcam | Clone TAL185, Cat ab20181 |
| Biological Samples |  |  |
| Tumor tissue | NKI | approved by the Institutional Review Board of the NKI CFMPB576 / IRB21-145 |
| Critical Commercial Assays |  |  |
| DAKO-PD-L1 IHC 22C3 pharmDx | DAKO | N/A |
| Opal 6-Plex Detection kit | Akoya Biosciences | Cat. NEL871001KT |
| AllPrep DNA/RNA/miRNA Universal Isolation Kit | Qiagen | 80224 |
| AllPrep DNA/RNA FFPE Isolation Kit | Qiagen | 80234 |
| QIAamp DNA formalin-fixed paraffin-embedded Tissue Kit | Qiagen |  |
| Software and Algorithms |  |  |
| HALO | Indica Labs | v3.2 |
| HALO HighPlex FL | Indica Labs | v4.0.2. |
| Slidescore | Internet | www.slidescore.com |
| InForm Software | Inform software | version 2.5.0 |
| BWA aln |  | version 0.7.17 |
| qDNAseq package |  | v1.12 |
| Other |  |  |
| L-PAC |  | https://github.com/dmmiedema2/LPAC/ |

#### Patient selection and sample collection

A total of 170 tumor samples from 24 EAC patients with initially locally advanced disease, who developed tumor recurrence after being treated with nCRT and surgical resection in the Netherlands Cancer Institute (NKI), were included in the study. The samples were collected at the following time points: before nCRT (*preCRT*), after nCRT (*postCRT*) and at *recurrence* of disease (Figure 1A). Clinical and pathological data were retrieved from the electronic patient files. The study was approved by the Institutional Review Board of the NKI (CFMPB576 / IRB21-145).

#### DNA Isolation

DNA was isolated from formalin fixed paraffin embedded (FFPE) tissue samples. Between 5 to 10 (depending on tumor size) 10 μm slides were used for isolation. DNA and RNA was isolated simultaneously with the AllPrep DNA/RNA FFPE isolation kit (Qiagen, #80234) by using the QIAcube, according to manufacturer’s protocol. Quantification and quality assessment of DNA were performed with a Nanodrop 2000 (Isogen), Qubit 2.0 Fluorometer (Invitrogen) and Tapestation 4200 (Agilent).

The tumor cell percentage was determined on haematoxicilin and eosine (H&E) stained slides, which were cut before and after the slides used for DNA/RNA isolation. The area’s with the highest tumor cell percentage were marked on these H&E slides by an experienced gastrointestinal pathologist (LK). The mean tumor cell percentage, determined on these two slides, was used for analysis.

#### DNA sequencing

Next-generation sequencing (NGS) library preparation were performed as described by Scheinin et al.(40). Briefly, 250ng genomic DNA was extracted with the QIAamp DNA formalin-fixed paraffin-embedded Tissue Kit (Qiagen, Hilden, Germany) and fragmented by ultrasonification (Covaris ME220; Covaris Inc., Woburn, MA). Sheared DNA was used for NGS library preparation with unique indexes (IDT, Coralville, IA) following the automated SOLiDTM procedures (Applied Biosytems, Foster City, CA, USA), followed by Sequencing with the HiSeq 4000 50bp single read (Illumina, San Diego, CA).

#### DNA copy number data pre-processing

Shallow whole-genome DNA sequencing (sWGS) reads were aligned using bwa aln version 0.7.17. Relative and segmented copy number profiles from the DNA sequencing reads and concurrent segmentation of these profiles was done using the qDNAseq package v1.12, using 1000kbp bins (40).

#### L-PAC multi-region inference of absolute CNAs

The relative genome-wide segmented copy numbers yielded by qDNAseq are used as input for L-PAC to infer absolute CNAs from multi-sample data. The multi-sample data processed by L-PAC should be genetically strongly related, e.g. acquired from the same cancer. Hence, multi-sample inference of CNAs by L-PAC is performed per patient in this study (Figure 1C). The procedure to infer absolute CNAs by L-PAC is as follows:

1. **Single sample inference**. For each sample from a tumor a separate estimate of the absolute CNAs is produced by the inference method we previously introduced (25). We repeat the description of the method here, with a generalization to account for noise. The measured relative copy number *r*_*i*_ of segment *i* with width *w*_*i*_ is determined by the absolute copy number *q*_*i*_ of, and the noise *η*_*i*_ on, segment *i* and tumor cell purity *α* of the sample:

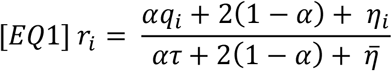 Where the tumor ploidy 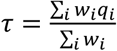 and the 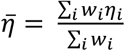 average noise. In the following we assume noise is random and hence averages to zero: 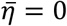 Eq. (1) can be rewritten to an expression for *q*_*i*_:

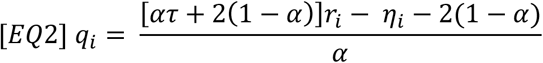

the quantity of interest to derive by inference from the measured *r*_*i*_’s. Note that noise *η*_*i*_ cannot be measured independently and is hence an unknown in Eq. (2). Inference proceeds by assuming that noise is zero, or at least negligible upon inference of absolute CNAs. A grid search over a range of purities *α’* = 0.2, 0.21, .., 1 and ploidies *α’* = 1.5, 1.55, .., 5 is performed to find the *q*_*i*_’s that best fit to integer values. The closest distance to an integer value *d*_*i*_ is defined as:

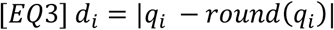

such that the best fit to integer values and hence absolute CNAs *q*_*i*_ are found by:

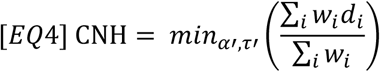 Where the minimal value of the grid search immediately defines the karyotype ITH: copy number heterogeneity (CNH). The rationale behind this inference method is that with sufficiently low noise and karyotype ITH, the measured *r*_*i*_ values are periodically spaced with the offset and periodicity defined by *α* and *α* (23). In that scenario, CNAs in the bulk sample value are close to integer values (i.e., *d*_*i*_ ≈ 0).
2. **Identification of failed inference**. We next explore what happens if the inference fails, which indeed is expected to happen when the assumptions of sufficiently low noise and low karyotype ITH are violated. We first note from Eq. (1) that the signal-to-noise ratio in the measured value *r* is 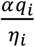 and hence decreases proportionally with tumor cell purity *α*. This implies that at low levels of *α, r*_*i*_ becomes dominated by stochastic noise (Eq. (1)). Because *η*_*i*_ cannot be measured independently and is assumed to be zero for inference, also *q*_*i*_ becomes dominated by stochastic noise (Eq. (2)). It follows that the minimization procedure in Eqs. (3 and 4) becomes a minimization of fluctuations around the lowest available integer value of *q*_*i*_ in the grid search (*α* ≈ 2), which is achieved at maximum *α*^*’*^(i.e. *α* = 1). Hence, failed inference can be recognized from the produced output by a purity = 1 and a ploidy < 2.5.
3. **Identification of best sample**. From the samples for which single-sample inference as performed in step 1) was successful according to the criterion in step 2), we identify the sample with the lowest karyotype ITH, defined by CNH, as the best sample β with the most reliable single-sample absolute CNA inference result.
4. **Alignment to CNAs of best sample**. For all but the best sample β a second grid search is performed over *α* and τ to infer absolute CNAs. The inference procedure is analogous to the procedure as described in step 1), with one crucial difference. In step 1) the inference yields the absolute CNA that best aligns to integer values (Eq. (3)). Here, the alignment is performed to the absolute CNAs of the best sample β, such that Eq. (3) in the inference procedure of step 1) is replaced with:

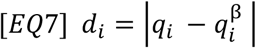 Where 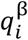 is the absolute CNA at segment *i* of the best sample β produced by single-sample inference. Please note that the underlying assumption is the sequential accumulation of CNAs affecting single chromosomes. For sub-clonal whole-genome doubling events the assumption behind L-PAC is invalid. Hence, for tumors or cancer types where sub-clonal whole genome doubling might be expected, the results from L-PAC should be interpreted with caution.

### CNA gains and losses

Gains and losses are defined from relative CNAs as the segment CNA value *r*_*i*_ > 1.1 or *r*_*i*_ < 0.9, respectively. For absolute CNAs, gains and losses are called if *q*_*i*_/*α* > 1.25 or *q*_*i*_/*α* < 0.75, respectively.

### Multiplex immunofluorescence analysis with multispectral imaging

Multiplex immunofluorescence (IF) analysis was performed on 98 formalin-fixed paraffin-embedded (FFPE) slides of the *postCRT* samples of each patient. A total of 118 regions, corresponding to areas selected for DNA isolation, were analyzed. Prior to multiplex staining 3μm slides were cut on TOMO slides. Slides were then dried overnight and stored in +4°C. Before a run was started slides were baked for 30 minutes at 70°C in an oven. Staining was performed on a Ventana Discovery Ultra automated stainer, using the Opal 6-Plex Detection Kit (50 slides kit, Akoya Biosciences, cat NEL871001KT). Protocol starts with baking for 28 minutes at 75°C, followed by dewaxing with Discovery Wash using the standard setting of 3 cycles of 8 minutes at 69°C. Pretreatment was performed with Discovery CC1 buffer for 64 minutes at 95°C, after which Discovery Inhibitor was applied for 8 minutes to block endogenous peroxidase activity. Two multiplex IF panels were used, MPIF21 and MPIF20, detecting specific markers consecutively on the same slide with the following antibodies. The first panel (MPIF21): Anti-CD68 (Clone KP1, Cat M0814, DAKO, 1/300 dilution 1h at RT), anti-CD8 (Clone C8/144B, Cat M7103, DAKO, 1/100 dilution, 1h at RT), anti-FoxP3 (clone 236A/E7, Cat ab20034, Abcam, 1/100 dilution, 2h at RT), anti-CD3 (Clone SP7, Cat RM-9107-S,Thermo Scientific, 1/400 dilution 1h at RT), Anti-PanCK (Clone AE1/AE3, cat MS343P, Thermo Scientific, 1/100 dilution, 2h at RT) and Anti-CD20 (Clone L26, Cat M0755, DAKO, 1/500 dilution, 1h at RT). The second panel (MPIF20): Anti-CD11b (Clone D6X1N, Cat CST49420s, Cell Signaling, 1/50 dilution 2h at RT), anti-CD3 (Clone SP7, Cat RM-9107-S,Thermo Scientific, 1/400 dilution 1h at RT), anti-CD15 (Clone MMA, Cat 559045, Becton Dickinson, 1/100 dilution, 1h at RT), anti-CD14 (clone EPR3653, Cat ab133335, Abcam, 1/100 dilution, 2h at RT), Anti-PanCK (Clone AE1/AE3, cat MS343P, Thermo Scientific, 1/100 dilution, 2h at RT) and Anti-HLA-DR (Clone TAL185, Cat ab20181, Abcam, 1/2000 dilution, 1h at RT). Each staining cycle was composed of four steps: Primary Antibody incubation, Opal polymer HRP Ms+Rb secondary antibody incubated for 1h at RT, OPAL dye incubation (OPAL480, OPAL520, OPAL570, OPAL620, OPAL690, OPAL780, 1/40 or 1/50 dilution as appropriate for 1 hour at RT) and an antibody denaturation step using CC2 buffer for 20 minutes at 95°C. Cycles were repeated for each new antibody to be stained. At the end of the protocol slides were incubated with DAPI (1/25 dilution in Reaction Buffer) for 12 minutes. After the run was finished slides were washed with demi water and mounted with Fluoromount-G (Southern Biotech, cat 0100-01) mounting medium.

After staining slides were imaged using the Vectra Polaris automated imaging system (Akoya Biosciences). Scans were made with the MOTiF protocol. Using the InForm software version 2.5.0 the MOTIF images were unmixed into 8 channels: DAPI, OPAL480, OPAL520, OPAL570, OPAL620, OPAL690, OPAL780 and Auto Fluorescence. In the MPIF21 panel the channels consisted of: DAPI, Pan-cytokeratin (PanCK) (Opal 480), CD3 (Opal 520), CD8 (Opal 690), FoxP3 (Opal 570) CD68 (Opal 620), CD20 (Opal 780) and auto fluorescence. The MPIF20 panel consisted of: DAPI, PanCK (Opal 480), CD15 (Opal 520), CD11b (Opal 570), CD14 (Opal 620), CD3 (Opal 690), HLA-DR (Opal 780) and auto fluorescence. Both were exported to a multilayered TIFF file.

The multilayered TIFFs were fused with HALO software version 3.2 Indica Labs to create one file for each sample.

The Indica Labs’ HALO HighPlex FL v4.0.2 analysis algorithm was used for the analysis using AI nuclei segmentation. The MPIF20 panel was used to identify myeloid derived suppressor cells (MDSC) (41). We phenotyped CD11b^+^CD14^−^CD15^+^ cells as granulocytic/polymorphonuclear MDSCs (PMN-MDSCs) and CD14^+^CD15^−^HLA-DR^lo/–^ cells as monocytic MDSCs (M-MDSCs) (41).

A classifier was created for each panel to calculate the immune cell density in stromal and intra-epithelial areas, expressed as immune cells/mm2.

### PD-L1 staining and assessment of PD-L1 expression

PD-L1 staining was performed using the DAKO-PD-L1 IHC 22C3 pharmDx on a BenchMark Ultra autostainer (Ventana Medical Systems) on FFPE samples. Samples were considered evaluable if at least ≥100 viable tumor cells were present. CPS was calculated as the number of tumor cells with PD-L1 cell membrane staining plus the number of immune cells (lymphocytes and macrophages) with cell membrane or intracellular PD-L1 staining, divided by the total number of viable tumor cells, multiplied by 100. Scoring was performed by an experienced gastro-intestinal pathologist (L.K.).

### Analysis of iTME

The iTME was characterized as described above (Section: *Multiplex immunofluorescence analysis with multispectral imaging)* for 100 *postCRT* regions from 18 tumors in which inference by L-PAC was successful and which had measurements from more than one *postCRT* region. The iTME of each region was studied in relation to the genomic similarity of that region compared to the sample of tumor recurrence. The similarity to recurrence of a *postCRT* sample was defined as the inverse of dCNA between the *postCRT* sample and the recurrence, divided by the average inverse dCNA of all *postCRT* samples of the tumor. The normalization per tumor was performed to reduce inter-tumor variations in combined analysis of data from all EACs. Similarly, the inter-tumor variation in the iTME was corrected for by dividing cell type density of a region by the average density of all regions measured in a tumor.

### Statistical analysis

Baseline characteristics are presented as mean (Standard Deviation (SD)) for a normally distributed variables or median (interquartile range (IQR)) for non-normally distributed variables and as counts (percentage) for categorical variables. Statistical tests used are reported in the text at the appropriate place. Matlab R2020a was used for data analysis.

