## Supplementary material for "Karyotype Evolution in Response to Chemoradiotherapy and Upon Recurrence of Esophageal Adenocarcinomas"

| | Total patients: n=24 | | Total analyzed patients: n=20 | |
|---|---|---|---|---|
| | n | % | n | % |
| **Age** *(median (IQR))* | 62 (51.3-65) | | 62 (52.5-65) | |
| **Male** | 23 | *96%* | 19 | *95%* |
| **Adenocarcinoma** | 24 | *100%* | 20 | *100%* |
| **Differentiation grade** | | | | *0%* |
| *Well/moderately differentiated* | 16 | *67%* | 16 | *80%* |
| *Poorly differentiated* | 8 | *33%* | 4 | *20%* |
| **Tumor location** | | | | *0%* |
| *Mid-esophageal* | 1 | *4%* | 1 | *5%* |
| *Distal esophagus* | 16 | *67%* | 12 | *60%* |
| *Esophago-gastric junction* | 7 | *29%* | 7 | *35%* |
| **Stage of disease at diagnosis*** | | | | *0%* |
| *Stage 3* | 16 | *67%* | 13 | *65%* |
| *Stage 4* | 8 | *33%* | 7 | *35%* |
| **Smoking** ***yes*** | 13 | *54%* | 10 | *50%* |
| **Alcohol** ***yes*** | 15 | *63%* | 13 | *65%* |
| **Radiotherapy dose** | | | | *0%* |
| *36* | 2 | *8%* | 1 | *5%* |
| *41,4* | 17 | *71%* | 15 | *75%* |
| *45* | 1 | *4%* | 1 | *5%* |
| *50,4* | 4 | *17%* | 3 | *15%* |
| **Chemotherapy** | | | | *0%* |
| carboplatin/paclitaxel | 18 | *75%* | 16 | *80%* |
| 5FU/cisplatin | 6 | *25%* | 4 | *20%* |
| **Stage of disease after nCRT + resection*** | | | | *0%* |
| *Stage 1* | 3 | *13%* | 2 | *10%* |
| *Stage 2* | 5 | *21%* | 5 | *25%* |
| *Stage 3* | 11 | *46%* | 9 | *45%* |
| *Stage 4* | 5 | *21%* | 4 | *20%* |
| **Mandard tumor regression grade (TRG)** | | | | *0%* |
| 1 | **1 | *4%* | 0 | *0%* |
| *2* | 7 | *29%* | 6 | *30%* |
| *3* | 9 | *38%* | 8 | *40%* |
| *4* | 7 | *29%* | 6 | *30%* |
| **Deceased** | 24 | 100% | 20 | *100%* |
| *Following disease progression* | 23 | 96% | 20 | *100%* |

** According to the 8th edition of the AJCC Cancer Staging Manual*
*** Patient with complete pathological response in CRT field, however with a satellite lesion which was analyzed*

**Table S1**
Clinical and pathological characteristics of included patients. AJCC = American Joint Committee on Cancer, nCRT= neo-adjuvant chemoradiotherapy, IQR= inter-quartile range

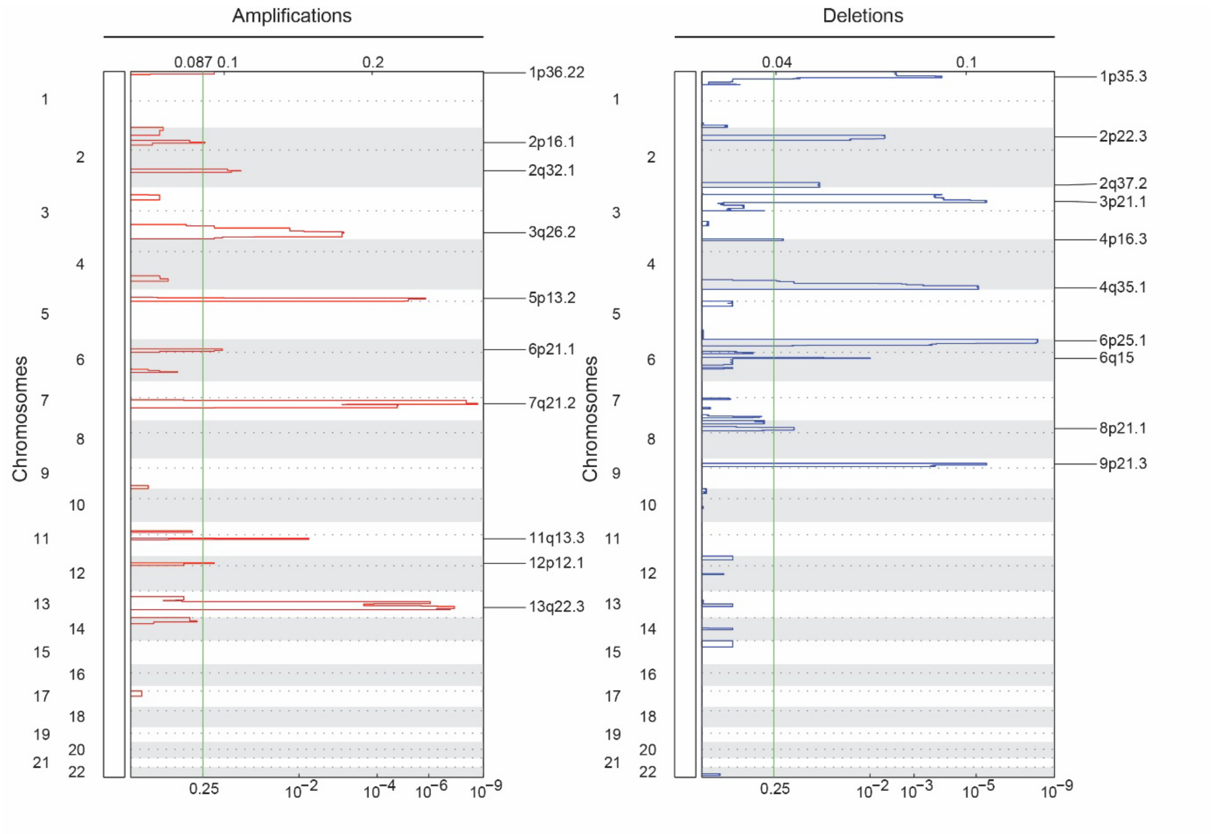


**Figure S1**

GISTIC analysis reveals recurrent focal CNAs in EAC.

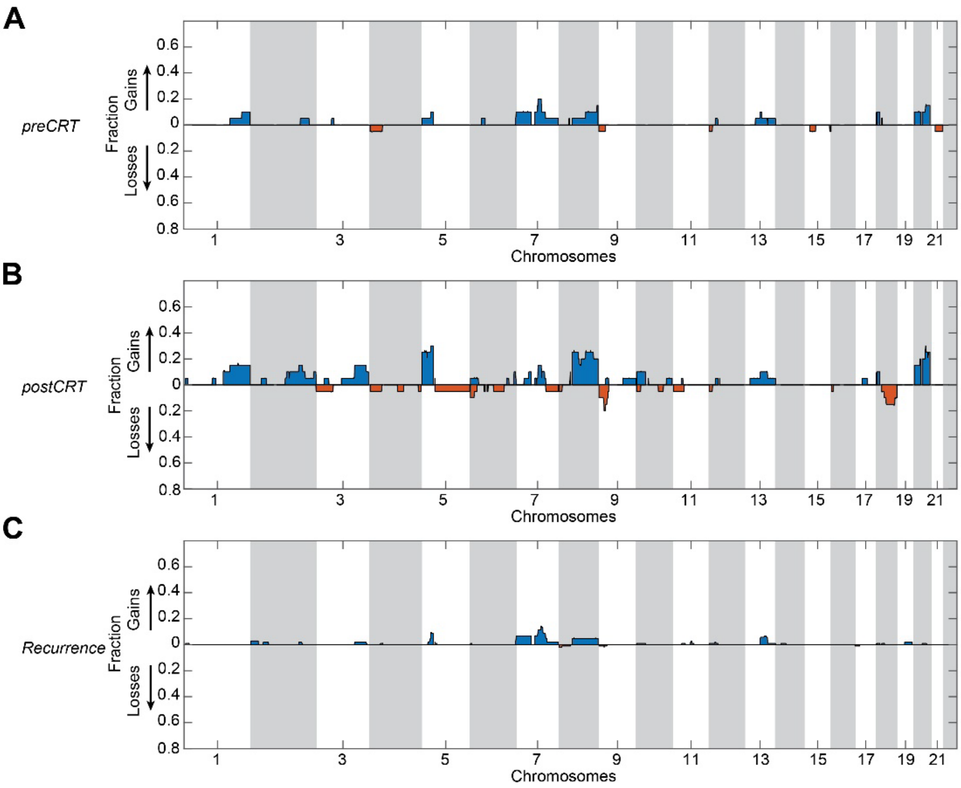


**Figure S2**

Fraction of gains and losses in absolute CNAs obtained using ABSOLUTE (23). **A,** Fraction gains and losses in *preCRT* samples. **B,** Fraction gains and losses in *postCRT* samples. **C, F**raction gains and losses in *recurrence* samples.

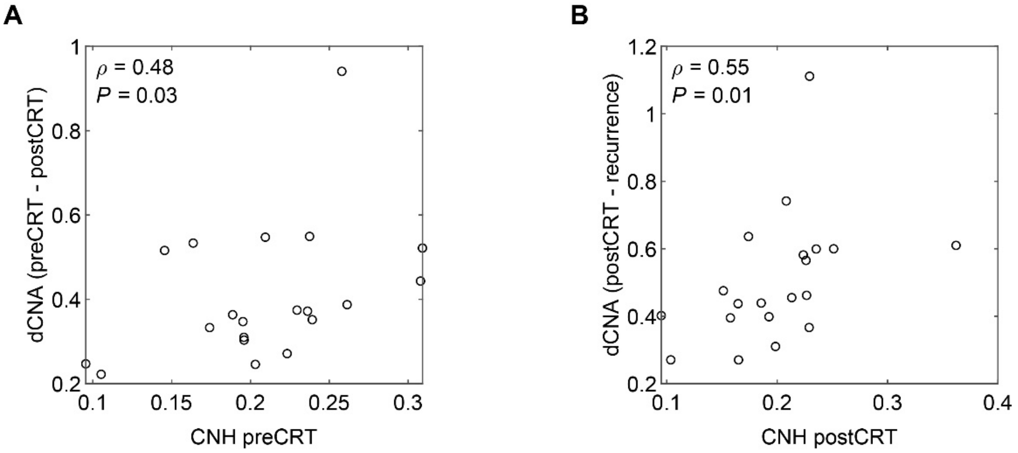


**Figure S3**

Prior level of copy number heterogeneity correlates with amount of karyotype evolution. **A,** dCNA in response to treatment versus copy number heterogeneity (CNH) in the *preCRT* sample. **B**, dCNA upon *recurrence* versus the average copy number heterogeneity (CNH) in the *postCRT* samples. The Spearman's rank correlation coefficients with corresponding p values are reported.

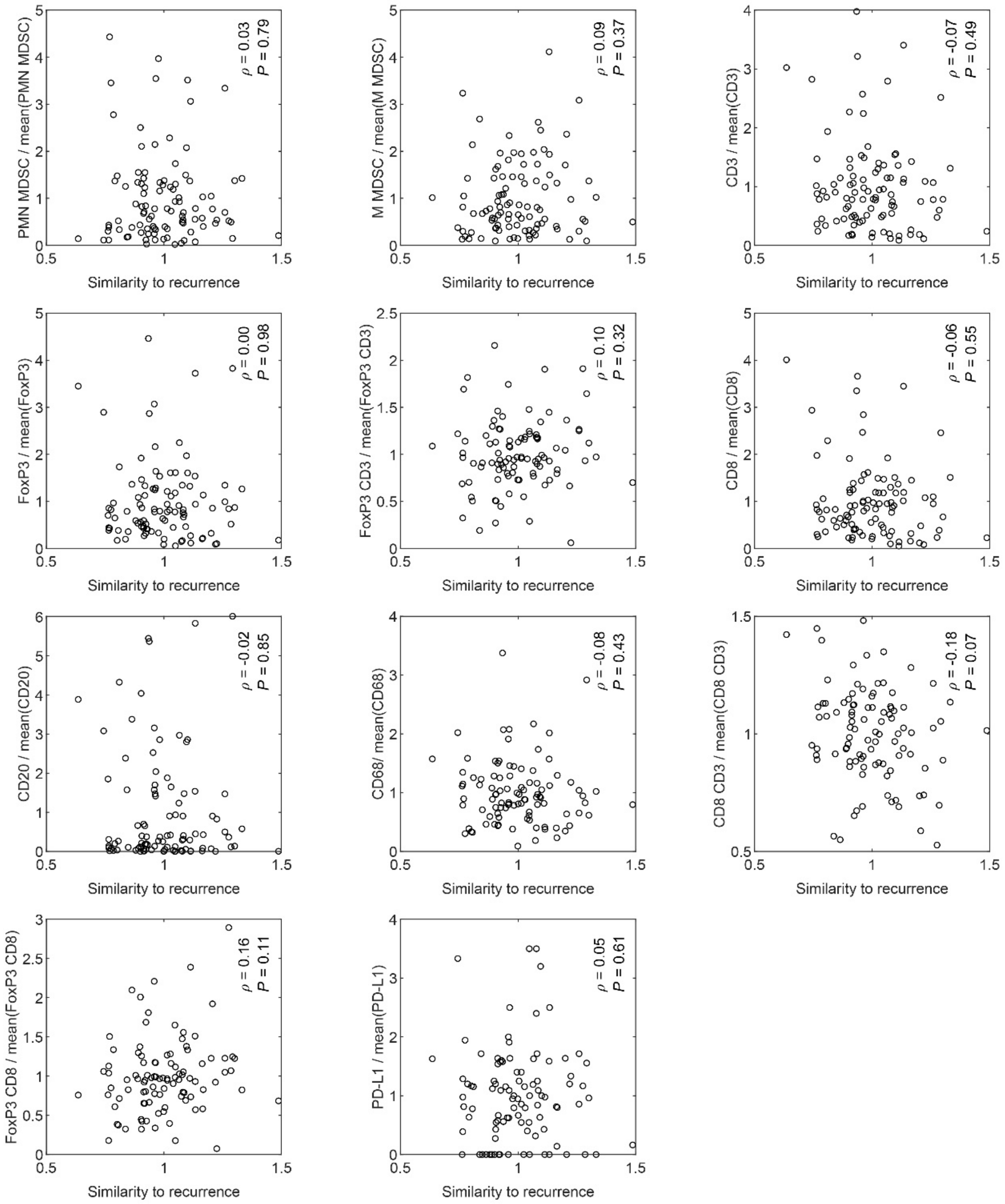


**Figure S4**

No correlation of iTME and similarity to *recurrence* of *postCRT* samples in combined analysis of patients with early and late recurrence. The Spearman's rank correlation coefficients with corresponding p values are reported.
